# SARS-COV-2 induces blood-brain barrier and choroid plexus barrier impairments and vascular inflammation in mice

**DOI:** 10.1101/2024.02.09.579589

**Authors:** Haowen Qiao, Xiangxue Deng, Lingxi Qiu, Yafei Qu, Yuanpu Chiu, Feixiang Chen, Shangzhou Xia, Cheyene Muenzel, Tenghuan Ge, Pengfei Song, Alexandre Bonnin, Zhen Zhao, Weiming Yuan

## Abstract

The coronavirus disease of 2019 (COVID-19) pandemic that has led to more than 700 million confirmed cases and near 7 million deaths. Although Severe Acute Respiratory Syndrome Coronavirus-2 (SARS-CoV-2) virus mainly infects the respiratory system, neurological complications are widely reported in both acute infection and long-COVID cases. Despite the success of vaccines and antiviral treatments, neuroinvasiveness of SARS-CoV-2 remains as an important question, which is also centered on the mystery whether the virus is capable of breaching the barriers into the central nervous system. By studying the K18-hACE2 infection model, we observed clear evidence of microvascular damage and breakdown of the blood-brain barrier (BBB). Mechanistically, SARS-CoV-2 infection caused pericyte damage, tight junction loss, endothelial activation and vascular inflammation, which together drive microvascular injury and BBB impairment. In addition, the blood-cerebrospinal fluid barrier at the choroid plexus was also impaired after infection. Therefore, cerebrovascular and choroid plexus dysfunctions are important aspects of COVID-19 and may contribute to the neurological complications both acutely and in long COVID.

## INTRODUCTION

After 4 years and over 700 million confirmed cases, the world is finally recovering from the COVID-19 pandemic. Despite the availability of successful vaccines and reasonably effective antiviral treatments, there are still unsolved mysteries related to its wide-ranging impacts on the body. While SARS-CoV-2 primarily infects the respiratory system, it often causes complications in other organs, including the brain, heart, kidneys and liver^1^. An intriguing fact of SARS-CoV-2 is its link with neurological complications, which is surprisingly common in COVID patients based on retrospective studies^2^. Besides common symptoms of impaired olfaction and gustation^3^, 36.4% of patients are reported exhibiting more severe neurological abnormalities, including impaired consciousness, acute cerebrovascular events, headache, etc^4^. The neurological symptoms may persist long after infection, and develop into the Post-acute Sequelae of COVID-19 (PASC)^5^. For example, delayed verbal memory, impaired immediate verbal memory and learning, depression and posttraumatic stress disorder symptoms were reported after 2 months following the infection^6^, and long-term cognitive dysfunctions were still seen at 7 months post-COVID^7^.

The blood–brain barrier (BBB), situated at the interface between the circulation and brain parenchyma, is specialized to prevent the invasion of parasites^8^, bacterial^9^, and viral pathogens^10,11^. The BBB comprises the brain microvascular endothelial cells, which are sealed together by the tight junctions and reinforced by extracellular basement membrane structures, perivascular mural cells, as well as astrocytic end feet^12^. Microvascular injuries including congested vessels and fibrinogen leakage were reported from post-mortem studies early in the pandemic^13^, while nearly one third of COVID-19 patients were later found to have high incidence of stroke^14^, venous thromboembolism^15^, intraventricular and subarachnoid hemorrhage^16^, and encephalitis^17^. Beside BBB, the brain is also protected by the blood-cerebrospinal fluid (CSF) barrier (BCSFB), which is located in the choroid plexus (ChP) tissues in the ventricles^18^. The ChP is highly vascularized, and its epithelial cells are the major producer of CSF in the brain, therefore it serves as a key interface separating the CSF from the blood^19^. It has been proposed that SARS-CoV-2 may entry the brain via the Angiotensin-converting enzyme 2 (ACE2) on the BBB^20^ or choroid plexus^21^. However, it is remains elusive if breakdown of the BBB occurs during infection, and whether cerebrovascular dysfunctions contribute to the neurological complications in COVID patients.

While studies from human post-mortem samples provided early evidence that SARS-CoV-2 has a huge impact on the cerebrovascular functions, in-depth mechanistic investigation in preclinical animal models remains limited. Although rodents are not natural hosts of SARS-CoV-2, transgenic mouse models carrying human ACE2 are highly susceptible to the virus and well-established for modeling severe COVID-19 disease^22^. Here using the K18-hACE2 model, we investigated the impact of SARS-CoV-2 infection on major parts of neurovascular systems, and found increased incidence of microhemorrhage and significant disruption of both BBB and BCSFB in K18-hACE2 mice after SARS-CoV-2 infection. Cerebral microvascular injury was accompanied by substantial pericyte damage, tight junction loss, astrogliosis and neuroinflammation in the brain parenchyma. In addition, endothelial activation and vascular inflammation occurred at both BBB and BCSFB, as shown by upregulation of VCAM-1 and COX2 markers. These results support that SARS-CoV-2 infection can directly induce microvascular and choroid plexus injuries, which may subsequently contribute to the neurological complications after infection and associated with PASC.

## MATERIALS AND METHODS

### Viruses, mouse strains and in vivo SARS-CoV-2 infection

The SARS-CoV-2 (Isolate USA-WA1/2020) was obtained from Biodefense and Emerging Infections Research Resources Repository (BEI Resources), propagated and titered in Vero-E6 cells as we previously described^23^, and was used for all in vivo studies. K18-hACE2-Tg (*B6.Cg-Tg(K18-ACE2)*^*2Prlmn/J*^) mice were purchased from the Jackson Laboratory (Stock #034860) and bred locally. K18-hACE2 transgene was maintained at hemizygous status in experimental mice. For SARS-CoV-2 infection, 1×10^4^ pfu virus were inoculated intranasally per mouse, as we recently reported^23^. Mice were monitored every day for weight loss and survival. All animal procedures were performed at ABSL3 facility at University of Southern California and approved by the Institutional Animal Care and Use Committee at the University of Southern California.

### Tissue collection and Histological analysis

After euthanasia, tissue samples were collected from mice, fixed in 4% paraformaldehyde (PFA) overnight. The tissues were embedded in paraffin and sectioned at 8-μm thickness at the USC Translational Pathology Core. For histology, paraffin sections were deparaffinized, rehydrated, then immersed in antigen retrieval solution (H-3300, Vector Laboratories) and submitted to heat-induced antigen retrieval. Then slides were washed with phosphate-buffered saline (PBS; pH 7.4), and blocked with 5% normal goat serum in PBST (0.2% Triton-X 100 in PBS) at room temperature for 30 min. The tissue sections were incubated with primary antibodies as indicated overnight at 4°C, followed by washing in PBST, and secondary antibody incubation for 1 hour at room temperature. Primary antibodies used in this study include: Claudin5 (1:800; Novus Biologicals: NB120-15106); ZO-1 (1:800; ThermoFisher Scientific: 40-2200); GFAP (1:500; ThermoFisher Scientific: 13-0300); CD13 (1:200; R&D System: AF2335); IBA1 (1:800; FujiFilm CDI R&D Sys: 019-19741); VCAM1 (1:800; EMD Millipore: CBL1300); COX2 (1:200; Cayman Chemical: 160126); CD31(1:800; ThermoFisher Scientific: MA3105); DyLight 649 conjugated Lectin from *Lycopersicon Esculentum* (1:400; ThermoFisher Scientific: L32472); Donkey anti mouse IgG (1:600; ThermoFisher Scientific: A-21202). Secondary antibodies used in this study were all purchased from ThermoFisher Scientific, which include Alexa Fluor 568 donkey anti-rabbit A10042; Alexa Fluor 568 donkey anti-rat IgG(H+L) A78946; Alexa Fluor 488 donkey anti-rabbit IgG(H+L) A21206; Alexa Fluor 488 donkey anti-mouse IgG(H+L) A21202; Alexa Fluor 568 donkey anti-mouse IgG(H+L) A10037.

For immunohistochemistry, ImmPRESS Duet Double Staining HRP/AP Polymer Kits (MP-7724 and MP-7724, Vector Laboratories) were used. Endogenous peroxidase activity was blocked, sections were incubated with Dual Specificity (anti-mouse IgG/rabbit IgG) ImmPRESS HRP/AP Polymer Reagent, following the manufacturer’s instructions. For substrate development, sections were incubated with ImmPACT DAB EqV Substrate (HRP, brown) for 5-10 minutes based on antigens. To ensure comparability, reactions for the same antibody staining were performed at the same time, with the same batch of DAB, with equal exposure to DAB for the same amount of time. After staining, sections were counterstained with hematoxylin (H-3502, Vector Laboratories), and mounted. For immunofluorescence staining, sections were blocked with 5% donkey serum in PBST. After primary antibody incubation overnight at 4°C, sections were stained with fluorescently conjugated secondary antibodies raised in Donkeys from ThermoFisher. Hoechst 33342 or DAPI (4’,6-diamidino-2-phenylindole) dye (1:5000) was used to mark cell nuclei.

Images were captured on a NikonTi2 confocal microscope equipped with an automated stage. For immunohistochemistry, the images were taken with a color camera. Low magnification images were obtained with tile scans and automatic stitching in the NIS-Elements Software. For immunofluorescence, images were taken with a confocal module, with the same laser intensity, detector sensitivity, amplification value, and offset were used for a given antibody staining. All images were analyzed with ImageJ software.

### Quantifications

The number of microhemorrhages was counted in the cortical area of each section. 3-5 nonadjacent sections per mice were used, and 10 randomized fields were imaged and analyze per mouse. For IgG leakage, the total number of blood vessels within the field of view and the number of vessels with IgG extravasation towards parenchymal were counted. The area occupied by the IgG signals were also calculated using the thresholding method in ImageJ. The levels of Claudin5, ZO-1 and CD13 were calculated based on integrated fluorescence intensity in ImageJ as we previously reported^24^, and normalized to the control group,

For endothelial VCAM1 and COX2 quantification, the VCAM1 or COX2 occupied vascular areas were calculated using the thresholding method in ImageJ, then compared with the Lectin or CD31 areas. For microglia and astrocyte assessment, the expression of IBA1 and GFAP were calculated using quantitative density measures, as we recently reported^25^.

### Statistical analysis

Descriptive statistics, including means and standard deviation (s.d.), were computed for each group. Data were presented as mean ± s.d. unless otherwise indicated. The sample size for each experiment is included in the figure legends. Statistical analyses were performed with two-tailed Student’s t-test as indicated in figure legends. *P < 0.05, **P < 0.01, ***P < 0.001 and ****P < 0.0001. NS denotes not significant (P value >0.05). All analyses were performed with GraphPad Prism (GraphPad Software).

## RESULTS

### SARS-CoV-2 infection in K18-hACE2 model

We infected the K18-hACE2 transgenic mice with the SARS-CoV-2 (Isolate USA-WA1/2020), at 1×10^4^ pfu per mouse via intranasal inoculation (**Fig. 1A**). Consistent with literature and as we recently reported, these mice developed severe symptoms after infection, including progressive weight loss (**Fig. 1B**) over the course of 7 days, and all mice died by 7 days post-infection or reaching euthanasia criteria (**Fig. 1C**). Next, we collected brain tissues at 5 dpi from infected mice. With immunohistochemistry, we found substantial amount of immunoreactivity to SARS-CoV-2 nucleocapsid (N) protein in the K18-hACE2 mouse brain (**Fig. 1D**), spreading throughout different brain regions including the cortex and thalamus. The presence of viral nucleocapsid protein indicates that active viral replication may have occurred in these areas. This is highly consistent with current evidence^26^ that SARS-CoV-2 virus can enter the brain in severe infection cases or in hACE2 models, which is presumably assisted by the hACE2 receptor at the brain endothelium^20^.

**Figure 1:**
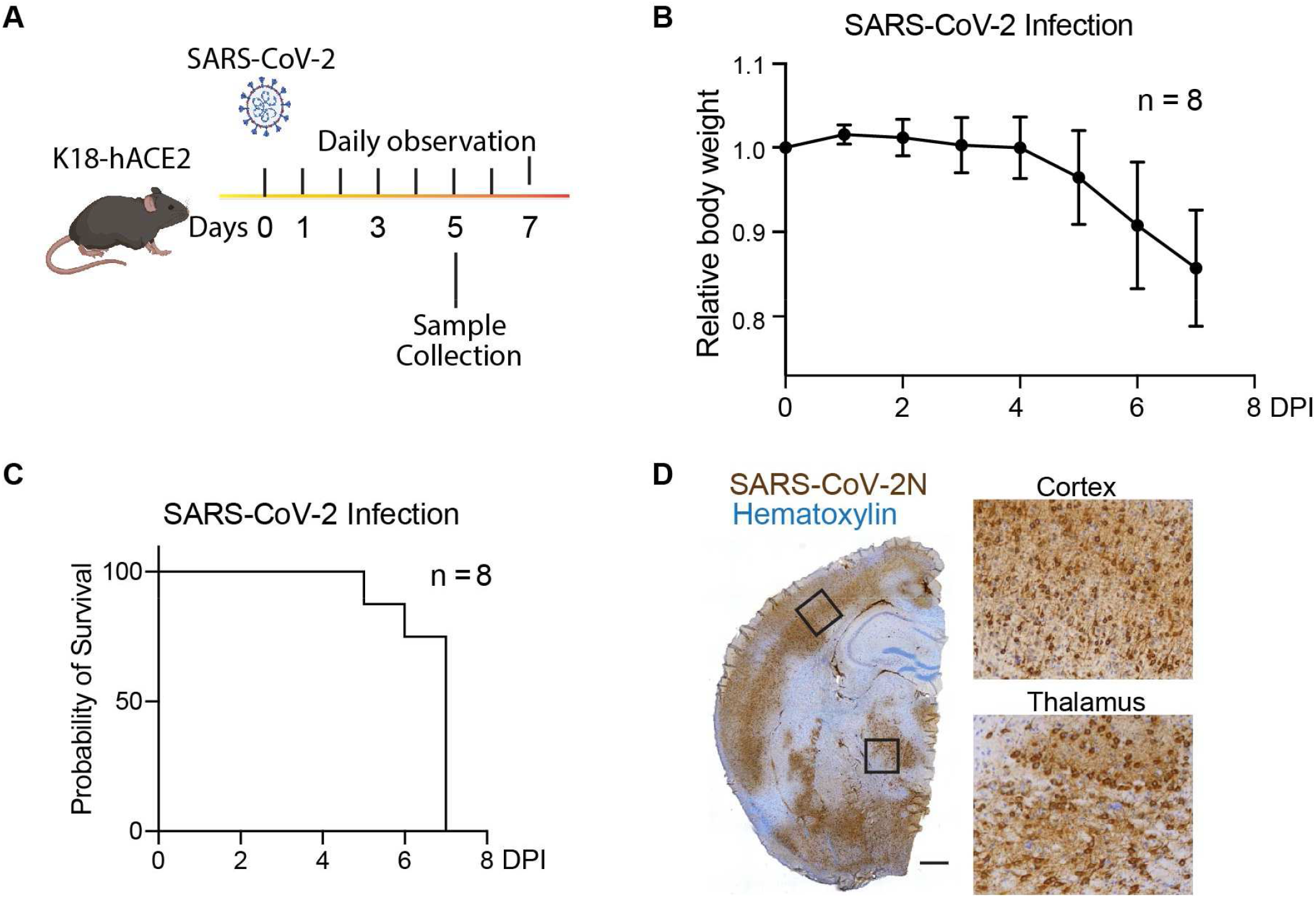
SARS-CoV-2 infection in the K18-hACE2 model. (**A**) Experimental diagram: K18-hACE2 mice were intranasally infected with SARS-CoV-2, monitored daily for symptoms, or euthanized at 5 days post-infection (DPI) to collect tissue. (**B**) The survival curve showing the probability of survival over 7 days after SARS-CoV-2 infection. (**C**) Daily weight measurements in K18-hACE2 mice intranasally infected with SARS-CoV-2 isolate USA-WA1/2020. (**D**) Representative mouse brain hemisphere image showing the presence of SARS-CoV-2 nucleocapsid protein by immunohistochemical staining. Boxed regions are shown on the right. Bar: 500 μm.

### Vascular damage and BBB breakdown in SARS-CoV-2 infected K18-hACE2 mice

To determine the vascular damage and potential BBB breakdown, we stained the sections for endogenous immunoglobin (IgG), and found it is not only accumulated in the microvessels but also reached the brain parenchymal (**Fig. 2A**). Next, we performed immunofluorescence staining, and found there were significantly increased incidence of microhemorrhage/microbleeds in the cortex and thalamus in the SARS-CoV-2 infected brains (**Fig. 2B-D**, Supplementary **Fig. S1A**). In the area without microhemorrhage, we still observe significantly increased IgG extravasation, including the cortex, thalamus, and hippocampus (Fig. **Fig. 2E-G**, Supplementary **Fig. S1B-C**). This was not due to the lack of perfusion after SARS-CoV-2 infection, as IgG in unperfumed mice was contained inside of the vessels (Supplementary **Fig. S1D**). These microvascular phenotypes are consistent with the postmortem studies throughout the pandemic, indicating that microvascular damages including congested vessels and BBB breakdown are integral to severe COVID-19^13^.

**Figure 2:**
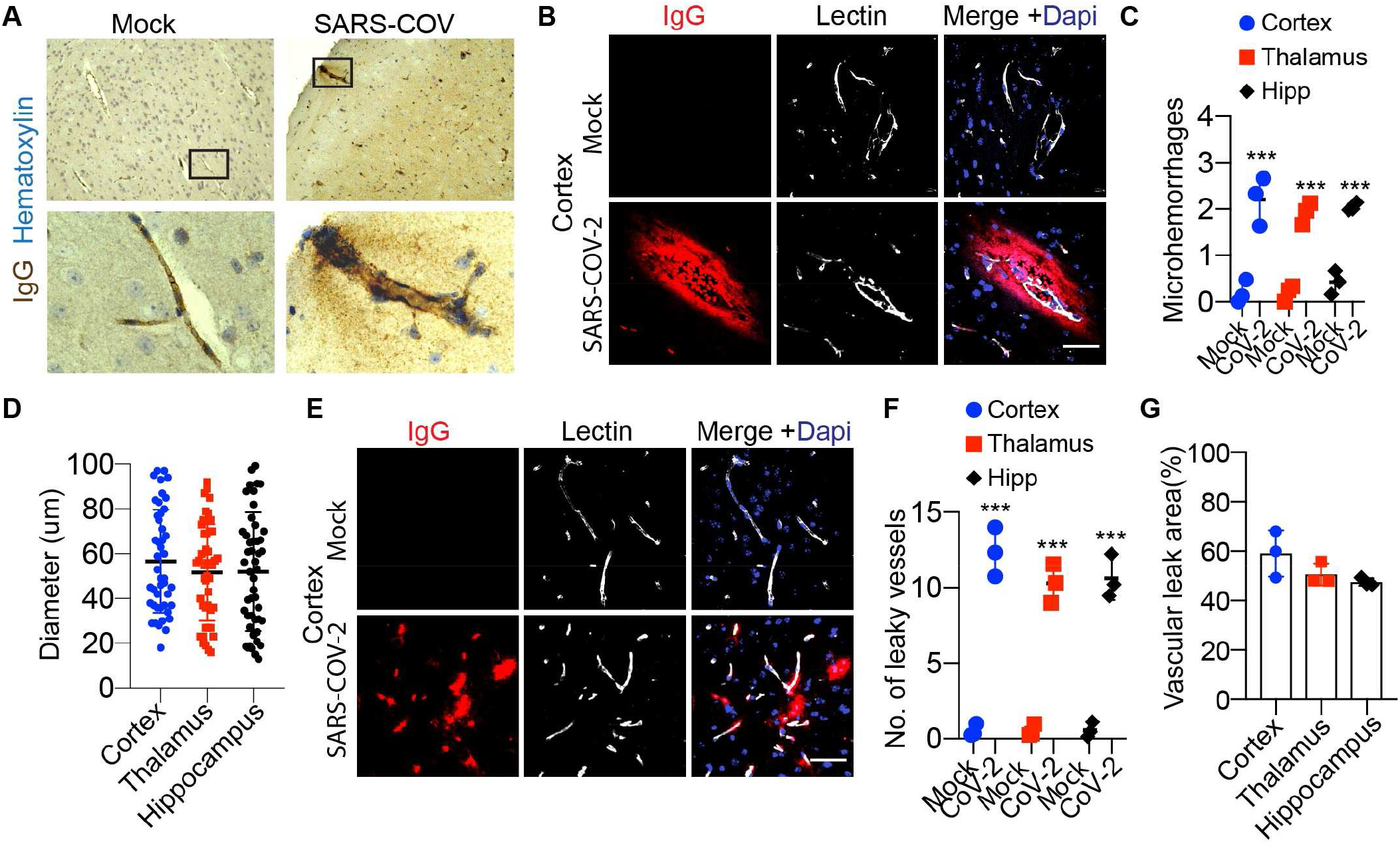
Vascular damage and BBB breakdown in SARS-CoV-2 infected K18-hACE2 model. (**A**) Representative images showing IgG immunohistochemical staining in K18-hACE2 mouse brain tissues with or without SARS-CoV-2 infection. Boxed regions are shown to the right. (**B**) Representative images showing immunofluorescent staining with IgG, showing the microhemorrhage site in the cortex of K18-hACE2 mice with SARS-CoV-2 infection. Bar: 50 μm. (**C**) Quantification of the numbers of microhemorrhages per field of view in cortex, thalamus, and hippocampus (Hipp). n= 3; ***p< 0.001; two-tailed Student’s t-test. (**D**) Quantification of the diameters of the microhemorrhages in the cortex, thalamus, and hippocampus. (**E**) Representative images showing immunofluorescent staining for IgG at the capillary level in the cortex. K18-hACE2 mice with SARS-CoV-2 infection exhibited significant accumulation of IgG surrounding the microvessels. Bar: 50 μm. (**F**) Quantification of the number of leaky small blood vessels per field of view in cortex, thalamus, and hippocampus. n= 3; ***p< 0.001; two-tailed Student’s t-test. (**G**) Quantification of percentage of area occupied by IgG per field of view in cortex, thalamus, and hippocampus. n= 3.

### The loss of tight junctions and pericytes in SARS-CoV-2 infected K18-hACE2 mice

Occludin, Claudins, and Zonula Occludens (also known as membrane-associated guanylate kinases, MAGUKs) are the major components of the tight junctions^27^, which preserve the polarity of the BBB and limit the entry of potentially neurotoxic plasma substances and pathogens into the brain. Here, we investigated the BBB structural changes using K18-hACE2 models upon SARS-CoV-2 infection (**Fig. 3A**). To verify the BBB integrity, we examined ZO-1 with immunostaining and found substantial losses of ZO-1-postive tight junctions in the cortex, thalamus, and hippocampus (**Fig. 3B-D**). Notably, the vascular segment with strong IgG extravasation tends to be the spot where tight junction protein ZO-1 were weakened (**Fig. 3B**). The relative fluorescence intensity of ZO-1 was decreased by more than 40% (**Fig. 3E**) in each brain area. In addition, the levels of Claudin5 in the Lectin-positive microvessels were also dramatically reduced in the cortex, thalamus, and hippocampus (**Fig. 3F-H**), by nearly 50% in each region (**Fig. 3I**).

**Figure 3:**
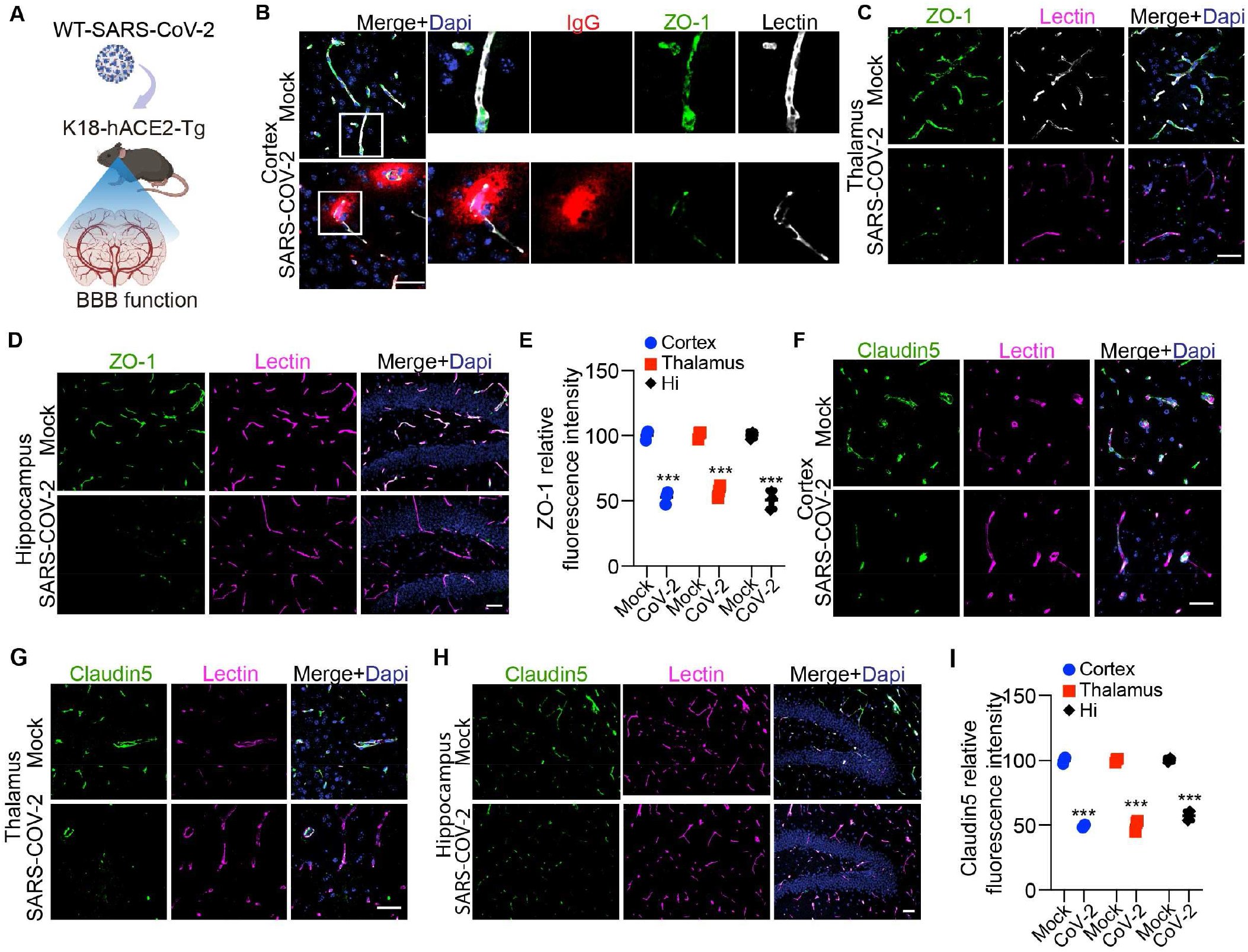
BBB tight junction loss in SARS-CoV-2 infected K18-hACE2 model. (**A**) The K18-hACE2 mice were used to test the BBB function after SARS-CoV-2 infection. (**B**) Representative images showing immunofluorescent staining for IgG, ZO-1, and Lectin in the cortex of K18-hACE2 mice. Bar: 50 μm. (**C-D**) Representative images showing immunofluorescent staining for ZO-1 and Lectin in the thalamus(**C**) and hippocampus(**D**) areas of K18-hACE2 mice. Bar: 50 μm. (**E**) Quantification of ZO-1 relative fluorescence intensity per field of view in cortex, thalamus, and hippocampus of K18-hACE2 mice. n= 3; ***p< 0.001; two-tailed Student’s t-test. (**F**-**H**) Representative images showing immunofluorescent staining for Claudin5 and Lectin in the cortex(**F**), thalamus (**G**), and hippocampus (**H**) areas of K18-hACE2 mice. Bar: 50 μm. (**I**) Quantification of the Claudin5 relative fluorescence intensity per field of view in cortex, thalamus and hippocampus. n= 3; ***p< 0.001; two-tailed Student’s t-test.

The extracellular basement membrane structures and perivascular mural cells are also components of the BBB, and substantially fortify the barrier. We next examined the brain pericytes using CD13 as a marker^28^, and found significant reductions of pericyte coverage along the microvessels in different brain regions, including cortex, thalamus, and hippocampus (**Fig. 4A-C**). The overall pericyte loss based on the CD13 levels are estimated to be over 40% in all the 3 major brain regions (**Fig. 4D**). These data demonstrate that the structural integrity of the BBB was severely impaired in K18-hACE2 mice after SARS-CoV-2 infection, including the loss of tight junctions and pericyte damage.

**Figure 4:**
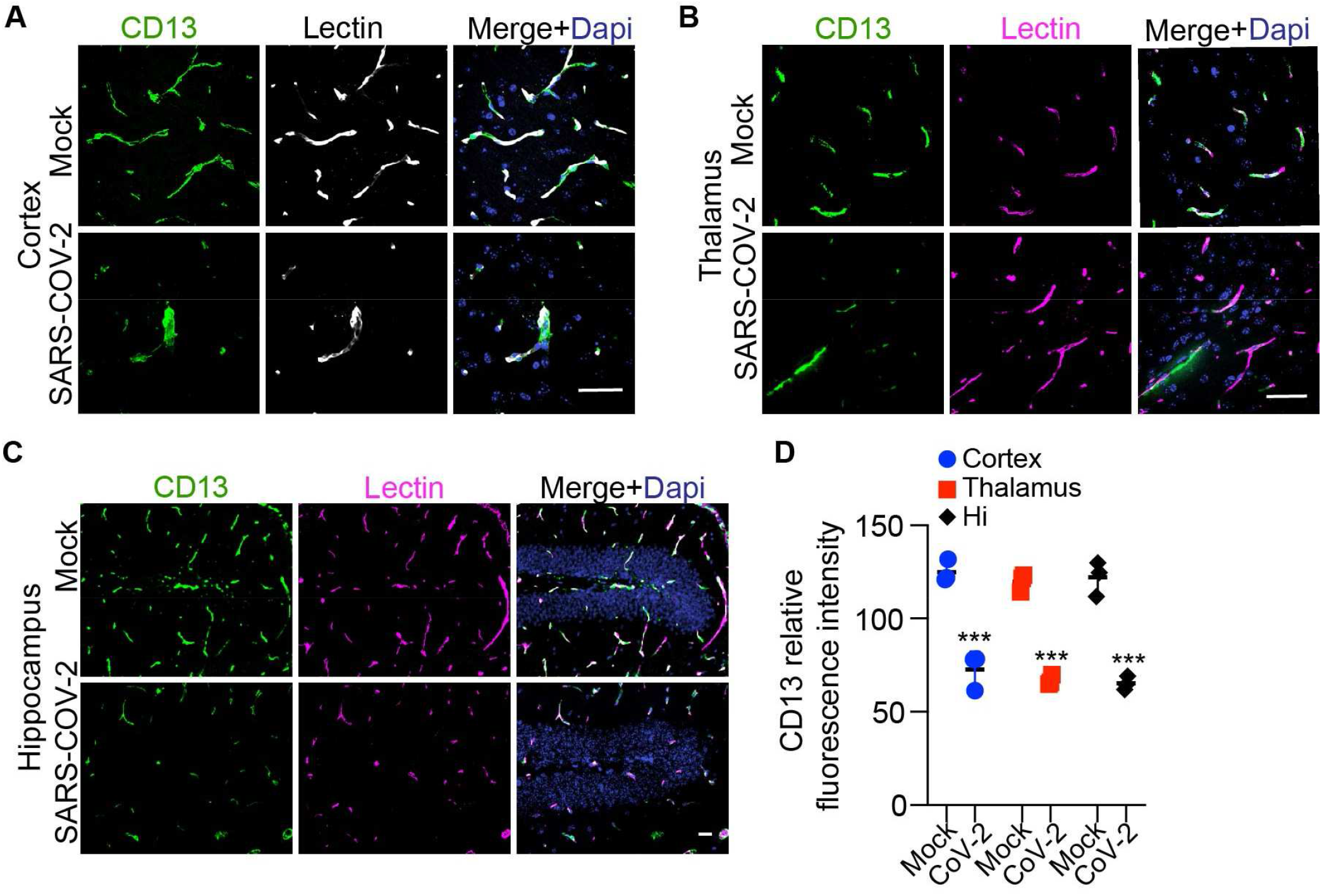
Loss of pericytes at the BBB of SARS-CoV-2 infected K18-hACE2 mice. (**A-C**) Representative images showing immunofluorescent staining for pericyte marker CD13 and endothelium marker Lectin in the cortex(**A**), thalamus (**B**), and hippocampus (**C**) areas of K18-hACE2 mice. Bar: 50 μm. (**D**) Quantification of the CD13 fluorescence intensity per field of view in the cortex, thalamus, and hippocampus. n= 3; ***p< 0.001; two-tailed Student’s t-test.

### Vascular inflammation and neuroinflammation in SARS-CoV-2 infected K18-hACE2 mice

Biomarkers studies have demonstrated significant upregulation of vascular biomarkers in COVID-19, including von Willebrand factor and C-reactive protein^29^. In addition, COVID-19 patients with neurological symptoms and multiple symptoms share three key biomarkers: TNFα, IL-6 and C-reactive protein^30^. Both TNFα and IL-6 are potent proinflammatory cytokines that stimulates vascular inflammation. Therefore, we examined endothelial activation and vascular inflammation using both vascular cell adhesion molecule 1 (VCAM-1) and cyclooxygenase-2 (COX2) markers. We observed a significant increase in vascular inflammation in BBB endothelial cells, as measured by VCAM-1 in lectin-positive microvessel profiles in cortex, thalamus, and hippocampus (**Fig. 5A-C**). While VCAM-1 is rarely expressed in microvessels in uninfected mice, it was found to be expressed in over 40% of the microvessels across different brain regions after SARS-CoV-2 infection (**Fig. 5D**).

**Figure 5:**
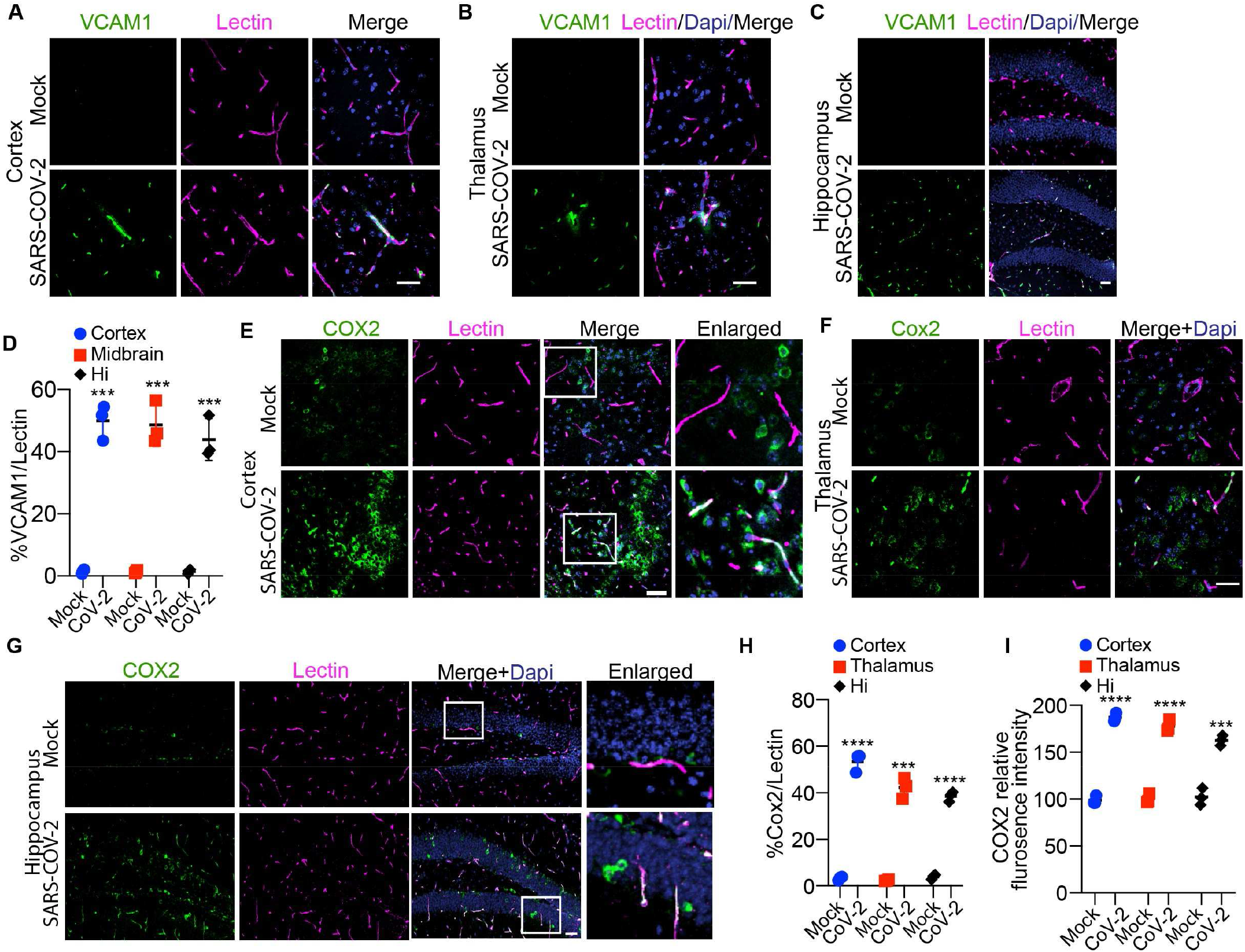
Vascular inflammation in SARS-CoV-2 infected K18-hACE2 model. (**A-C**) Representative images showing immunofluorescent staining for VCAM1 and Lectin in the cortex, thalamus, and hippocampus of K18-hACE2 mice. Bar: 50 μm. (**D**) Quantification of VCAM1 and lectin signal overlap in the cortex, thalamus, and hippocampus. n= 3; ***p< 0.001; two-tailed Student’s t-test. (**E-G**) Representative images showing immunostaining of COX2 and Lectin in the cortex, thalamus, and hippocampus of K18-hACE2 mice. Bar: 50 μm. (**J**) Representative images showing immunofluorescent staining for COX2 and CD31 in the ChP. (**H**) Quantification of COX2 and Lectin signal overlap in the cortex, thalamus, and hippocampus areas of K18-hACE2 mice. n= 3; ***p< 0.001; ****p< 0.0001; two-tailed Student’s t-test. (**I**) Quantification of COX2 relative fluorescence intensity in the cortex, thalamus, and hippocampus of K18-hACE2 mice. n= 3; ***p< 0.001; ****p< 0.0001; two-tailed Student’s t-test.

While COX2 is only minimally present in a subset of neurons in uninfected mice, SARS-CoV-2 infection increased COX2 levels dramatically in both neurons and cerebrovasculature (**Fig. 5E**). We also observed microvascular COX2 upregulation in the thalamus (**Fig. 5F**) and hippocampus (**Fig. 5G**). After measuring the proportion of COX2 signals in the lectin+ microvessel profiles, we found that over 40% of the cerebral vessels expressed COX2 after SARS-CoV-2 infection across all the three major brain regions (**Fig. 5H**), and the overall COX2 signals based on fluorescence intensity were also dramatically increased (**Fig. 5I**).

The upregulation of vascular biomarkers also coincides with the change of neuroinflammation status in the K18-hACE2 mouse brain after SARS-CoV-2 infection. We observed significant increases of astrogliosis in the cortex, thalamus, and hippocampus based on the GFAP marker (**Fig. 6A-C**), where GFAP-expressing astrocytes were nearly doubled (**Fig. 6D**). Substantial microglia activation also occurred in all three major brain regions after SARS-CoV-2 infection, as seen based on the IBA1 marker (**Fig. 6E-H**), indicting an interplay between the cerebrovascular and immune system.

**Figure 6:**
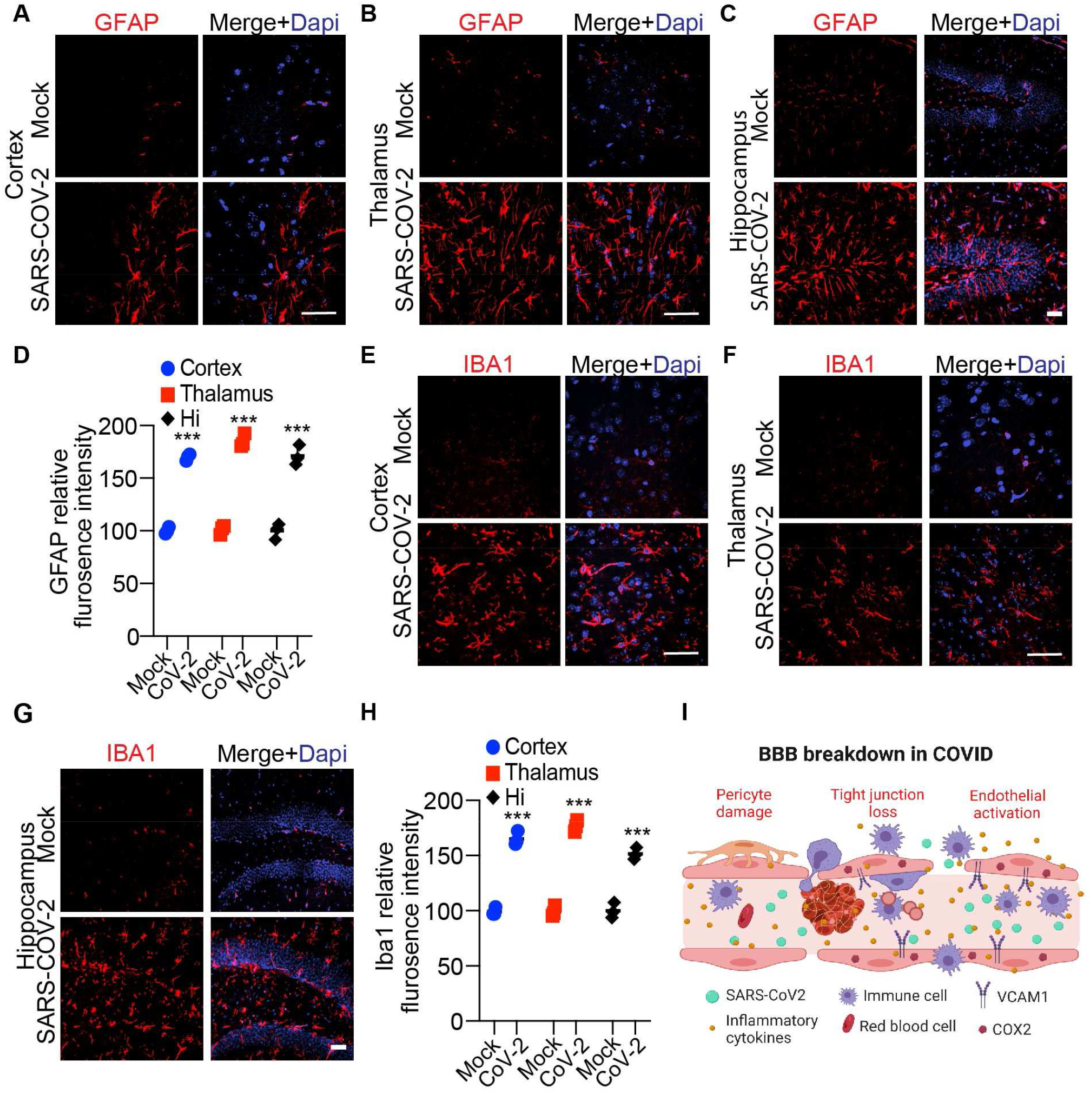
Astrogliosis and neuroinflammation in SARS-CoV-2 infected K18-hACE2 model. (**A**-**C**) Representative images showing immunofluorescent staining for GFAP in the cortex(**A**), thalamus (**B**), and hippocampus (**C**) of K18-hACE2 mice with or without SARS-CoV-2 infection. Bar: 50 μm. (**D**) Quantification of GFAP relative fluorescence intensity in the cortex, thalamus, and hippocampus of K18-hACE2 mice. n= 3; ***p< 0.001; two-tailed Student’s t-test. (**E**-**G**) Representative images showing immunofluorescent staining for IBA1 in the cortex (**E**), thalamus (**F**), and hippocampus (**G**) of K18-hACE2 mice. Bar: 50 μm. (**H**) Quantification of IBA1 relative fluorescence intensity in the cortex, thalamus, and hippocampus of K18-hACE2 mice. n= 3; ***p< 0.001; two-tailed Student’s t-test. (**J**) The diagram showing the major pathological events underlying the microvascular injury and BBB impairment in COVID-19. Severe SARS-CoV-2 infection led to pericyte loss, degradation of tight junctions, upregulation of VACM1 and COX2 molecules in the brain endothelium, attracting immune cells and causing local neuroinflammation. Virions also cross the BBB and replicate in the parenchymal and instigate more inflammatory responses and neuronal dysfunctions.

### Breakdown of the BCSFB in ChP in the SARS-CoV-2 infected K18-hACE2 mice

The BCSFB is also a critical interface between CNS and circulation, and we next tested whether the SARS-CoV-2 influenced the BCSFB function. We observed significant loss of epithelial tight junction structures in the ChP after SARS-CoV-2 infection, as indicated by both ZO-1 (**Fig. 7A-B**) and Claudin5 (**Fig. 7C-D**), indicating the breakdown of the BCSFB. In addition, we also found pericyte damage and loss based on CD13 staining (**Fig. 7E-F**), and substantial increases of both VCAM-1 and COX2 markers in the CD31-positive endothelium within the ChP after SARS-CoV-2 infection (**Fig. 7G-J**), as well as a strong activation of microglia within the ChP as shown by the IBA expression (**Fig. 7K-L**). This indicates that endothelial activation and vascular inflammation in ChP induced SARS-CoV-2 infection may influence the BCSFB functions.

**Figure 7:**
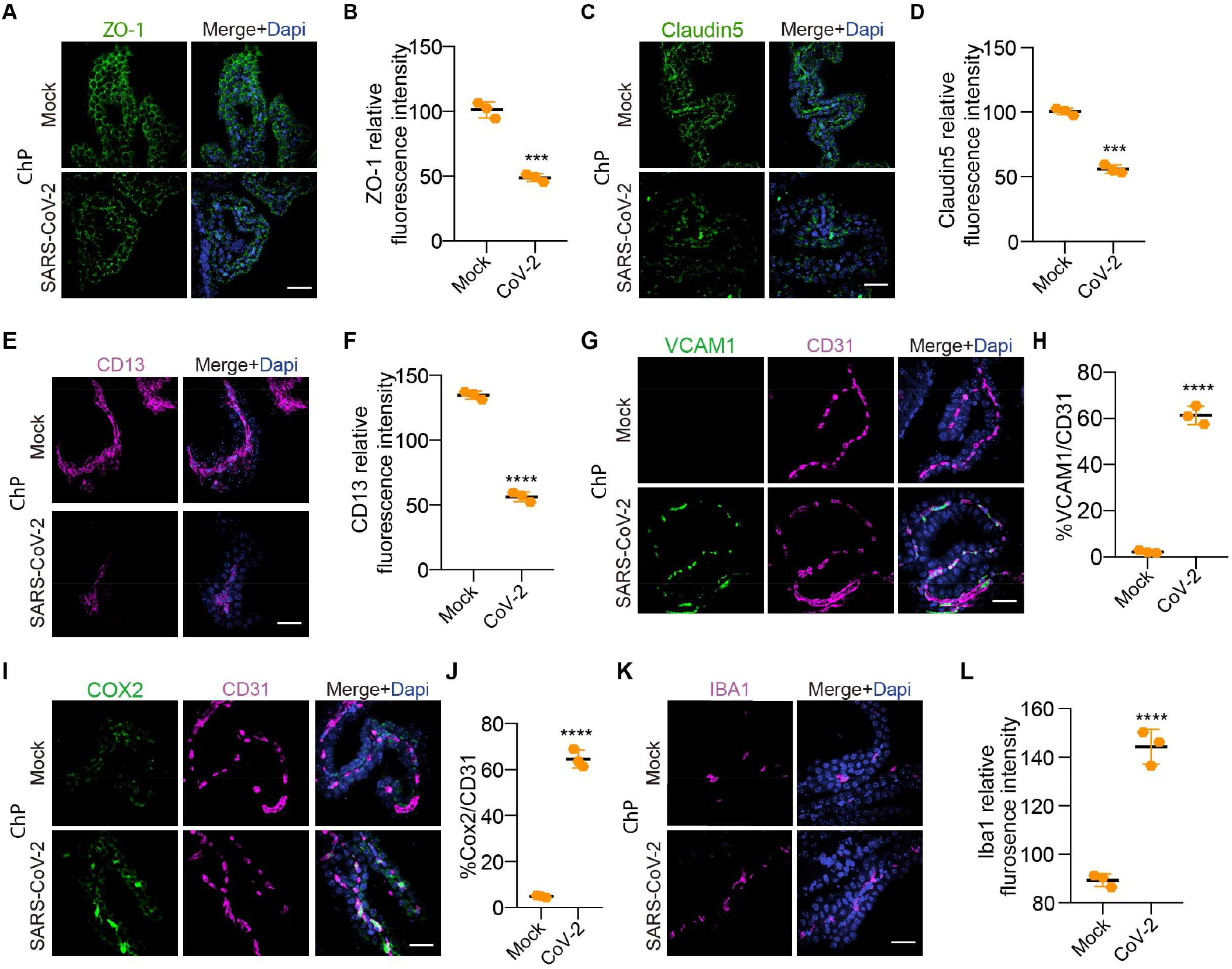
Breakdown of the BCSFB in ChP of the SARS-CoV-2 infected K18-hACE2 mice. (**A**) Representative images showing immunostaining of ZO-1 in the ChP of K18-hACE2 mice after SARS-CoV-2 infection. Bar: 50 μm. (**B**) Quantification of ZO-1 relative fluorescence intensity per field of view in ChP of K18-hACE2 mice. n= 3; ***p< 0.001; two-tailed Student’s t-test. (**C**) Staining for tight-junction protein Claudin5 in the ChP of K18-hACE2 mice after SARS-CoV-2 infection. Bar: 50 μm. (**D**) Quantification of Claudin5 expression in ChP of K18-hACE2 mice. n= 3; ***p< 0.001; two-tailed Student’s t-test. (**E**) Representative images showing immunostaining of CD13 in the ChP in the SARS-CoV-2-infected K18-hACE2 mice. Bar: 50 μm. (**F**) Quantification of CD13 expression in ChP of K18-hACE2 mice. n= 3; ****p< 0.0001; two-tailed Student’s t-test. (**G**) Staining for VCAM1 and CD31 in the ChP in the SARS-CoV-2-infected K18-hACE2 mice. Bar: 50 μm. (**H**) Quantification of VCAM1 and CD31 signal overlap in the ChP. n= 3; ****p< 0.0001; two-tailed Student’s t-test. (**I**) Representative image showing COX2 and CD31 in the ChP in the SARS-CoV-2-infected K18-hACE2 mice. Bar: 50 μm. (**J**) Quantification of COX2 and CD31 signal overlap in the ChP. n= 3; ****p< 0.0001; two-tailed Student’s t-test. (**K**) Representative images showing immunofluorescent staining for IBA1 in the ChP of K18-hACE2 mice. Bar: 50 μm. (**L**) Quantification of IBA1 relative fluorescence intensity in the ChP of K18-hACE2 mice. n= 3; ****p< 0.0001; two-tailed Student’s t-test

In sum, by studying the K18-hACE2 model with SARS-CoV-2 infection, we have observed clear evidence of microvascular damage, and breakdown of the BBB and BCSFB *in vivo*, demonstrating that the CNS barrier dysfunction is an important aspect of COVID-19. Mechanistically, pericyte damage, tight junction loss, endothelial activation, and vascular inflammation induced by SARS-CoV-2 infection may collectively drive cerebrovascular injuries and barrier impairments (**Fig. 6I**), which should be considered for intervention, particularly in preventing PASC or reducing the risk of developing it.

## Discussion

The central nervous system (CNS) is protected by the blood-brain barrier (BBB) and the blood-cerebrospinal fluid (CSF) barrier (BCSFB), yet both are potential routes and victims of SARS-CoV-2 virus invasion. Several studies supported that the BBB may be damaged in COVID patients. For example, Mathilde Bellon et al.^31^ presented the initial evidence of SARS-CoV-2-induced BBB disruption in 31 COVID-19 patients, and leakage was found in 58% of the cases. Additional clinical, laboratory, and autopsy evidence confirmed cerebrovascular impairments after SARS-CoV-2 infection, such as microvascular infarcts and hemorrhages, the prothrombotic or thrombophilic events, etc. Elevated D-dimer, a blood clotting marker, has been reported to be associated with COVID-19 severity in COVID-19 patients^32,33^. Like D-dimer, factor V is more active in patients with severe COVID-19, which associated with venous thromboembolism^34^. Neural imaging studies now confirmed patients with SARS-CoV-2 infection have much increased brain vascular permeability based on Dynamic contrast-enhanced magnetic resonance imaging (DCE-MRI)^35^. While all these evidence points to a critical role of cerebrovascular dysfunctions in the development of neurological manifestations associated with COVID-19, our basic understanding of the cellular and molecular changes in the brain vasculature remains limited. Thus, our study provided more clear evidence on the cerebrovascular injury and barrier dysfunctions post SARS-CoV-2 infection, and highlights the cellular and molecular events that drive the breakdown of the BBB and BCSFB, including tight junction loss, pericyte damage, endothelial activation, vascular inflammation. The impact of such changes, together with astrogliosis and neuroinflammation, may drive or at least contribute to the neurological complications seen in COVID-19 patients. As SARS-CoV-2 will likely remain as a major health issue for years to come, our findings provide a needed understanding of its impact on the major CNS barriers and brain homeostasis at both molecular and cellular levels.

Evidence from the hACE2 transgenic mouse model that is susceptible to SARS-CoV-2 infection indicated that SARS-CoV-2 may be able to pass the BBB with the help of angiotensin-converting enzyme 2 (ACE2) receptor, without causing significant damage to the BBB integrity^20^. Yet, human post-mortem studies^13^ and neural imaging follow-up in COVID-19 patients^35^, and *in vitro* cell models^36^ indicated that BBB dysfunction may be an important aspect. Mechanistically, Jan Wenzel et al. even reported that SARS-CoV-2 protease Mpro can cleave NEMO and impair nuclear factor-κB in brain endothelial cells^37^, and causing endothelial changes in mouse model of SARS-CoV-2 infection. Our study further indicated that the structural changes in cerebral microvessels and BBB are key pathological phenomenon during SARS-CoV-2 infection in the K18-hACE2 infection model. We reported a notable rise in the occurrence of microhemorrhages in the mouse brain infected with SARS-CoV-2. In areas without microhemorrhages, we also noted a substantial increase in IgG extravasation, particularly in the cortex, thalamus, and hippocampus. This underscores the significant impact of SARS-CoV-2 on microhemorrhages and emphasizes its impact on BBB integrity. Nevertheless, the current research remains limited on this direction, and future studies are still urgently needed to explore the fundamental biology in models with mild infection and long-COVID scenarios, and determine to what extent the breakdown of the brain barriers may contribute to the neurological complication in the development of PASC.

ChP cells also express ACE2, and is proposed as an alternative route of SARS-CoV-2 neuroinvasion^38^. This was further confirmed using ChP organoids, showing SARS-CoV-2 not only infected these organoids but also viral entered the ventricular CSF compartment^21^. Interestingly, Yang and colleagues^39^ conducted a study analyzing 65,309 single-nucleus transcriptomes using postmortem samples both from the frontal cortex and choroid plexus. Instead of finding molecular traces of SARS-CoV-2 in the cortex, they observed changes in the barrier cells of the choroid plexus and the infiltration of peripheral T cells into the parenchyma. Besides, it was also reported that the alterations in microglial and astrocyte subpopulations in COVID-19 patients were linked to the features commonly reported in human neurodegenerative diseases. However, it is still unclear that whether the SARS-CoV-2 infects ChP and impairs its barrier functions. By using the SARS-CoV-2 infected K18-hACE2 models, we found that SARS-CoV-2 also broke down the ChP epithelial barrier. The loss of tight junctions tightly correlated with pericyte damage, endothelial activation, and vascular inflammation in ChP *in vivo*, supporting the hypothesis that ChP dysfunction is an important aspect of neurological complications in COVID-19 patients.

SARS-CoV-2 infection induces extensive antiviral immune responses. Even though the CNS has immune privilege, neuroimmune responses could still occur after infection. However, the underlying mechanism of neuroimmune response post SARS-CoV-2 infection is very complex, including CD8+ T cell and microglia infiltration^40^, complement system activation^40,41^, IFN-I and proinflammatory signaling pathways initiation^42^. Marius Schwabenland et.al^43^ provided direct evidence of microglial and T cell infiltration in COVID-19 patients’ brain sections, furthermore, the microglia-T-cell interaction is tightly associated with neuroinflammatory parameters. It will be very interesting to further study the impact of infiltrating T cells and their potential roles in neuroinflammation. Our data additionally revealed broad astrogliosis and neuroinflammation in the cortex, thalamus, hippocampus, and ChP in SARS-CoV-2 infected K18-hACE2 models, suggesting that vascular impairment and BBB breakdown could contribute to the neuroimmune activation as well.

## Author contribution

H.Q., Z.Z. and W.Y. conceived of the research, designed the study, and wrote the manuscript. H.Q., X.D., L.Q., Y.Q., F.C., Y.C., S.X., C.M., and T.G. performed the experiments and analyzed data. W.Y. led the mouse infection experiments in BSL3 facility. P.S., and A.B. helped with the study design and manuscript writing. All authors commented on the manuscript.

## Acknowledgement

This work was supported by IDSA Foundation grant (Z.Z. and W.Y.). We thank Jill Henley, Dr. Lucio Comai and the Hastings and Wright Laboratories at USC for their support on the ABSL3 work.

---

**Supplementary Figure 1:**
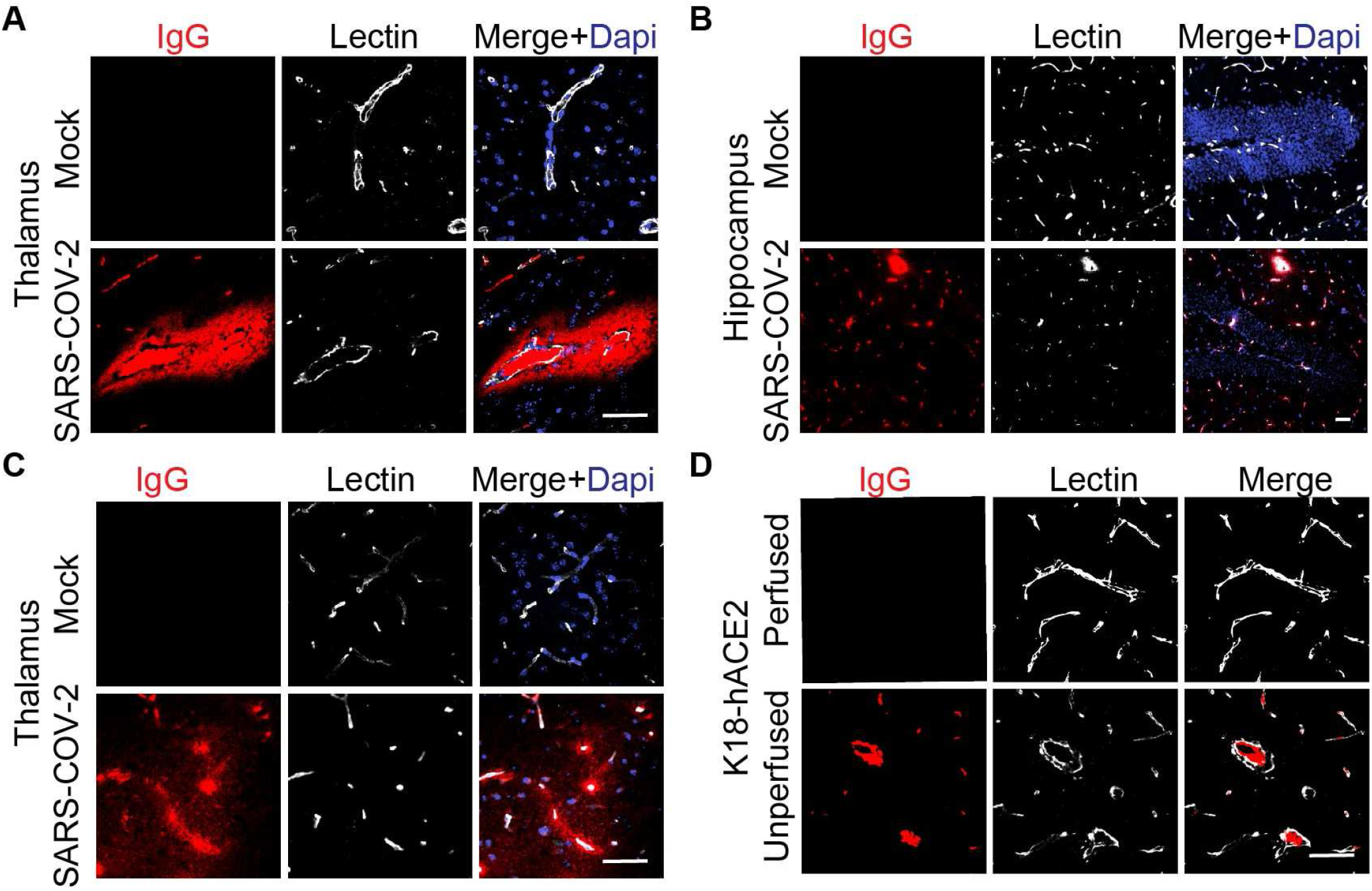
Additional data on vascular damage and potential BBB breakdown in SARS-CoV-2 infected K18-hACE2 model. (**A**) Representative images of immunofluorescent IgG staining showing microhemorrhage in the thalamus area of K18-hACE2 mice. Bar: 50 μm. (**B-C**) Representative images showing immunofluorescent staining for IgG at the capillary level in the hippocampus and thalamus areas. Bar: 50 μm. (**D**) Representative images showing IgG immunofluorescent staining in the cortex of perfused or unperfused mouse. Bar: 50 μm.

## References

1. Gavriatopoulou, M., Korompoki, E., Fotiou, D., Ntanasis-Stathopoulos, I., Psaltopoulou, T., Kastritis, E., Terpos, E., and Dimopoulos, M.A. (2020). Organ-specific manifestations of COVID-19 infection. Clin Exp Med 20, 493–506. 10.1007/s10238-020-00648-x.

2. Varatharaj, A., Thomas, N., Ellul, M.A., Davies, N.W.S., Pollak, T.A., Tenorio, E.L., Sultan, M., Easton, A., Breen, G., Zandi, M., et al. (2020). Neurological and neuropsychiatric complications of COVID-19 in 153 patients: a UK-wide surveillance study. The Lancet Psychiatry 7, 875–882. 10.1016/S2215-0366(20)30287-X.

3. Gupta, V., Banavara Rajanna, L., Upadhyay, K., Bhatia, R., Madhav Reddy, N., Malik, D., and Srivastava, A. (2021). Olfactory and Gustatory Dysfunction in COVID-19 Patients from Northern India: A Cross-Sectional Observational Study. Indian J Otolaryngol Head Neck Surg 73, 218–225. 10.1007/s12070-021-02391-5.

4. Mao, L., Jin, H., Wang, M., Hu, Y., Chen, S., He, Q., Chang, J., Hong, C., Zhou, Y., Wang, D., et al. (2020). Neurologic Manifestations of Hospitalized Patients With Coronavirus Disease 2019 in Wuhan, China. JAMA Neurol 77, 683. 10.1001/jamaneurol.2020.1127.

5. Munipalli, B., Seim, L., Dawson, N.L., Knight, D., and Dabrh, A.M.A. (2022). Post-acute sequelae of COVID-19 (PASC): a meta-narrative review of pathophysiology, prevalence, and management. SN Compr. Clin. Med. 4, 90. 10.1007/s42399-022-01167-4.

6. Elmazny, A., Magdy, R., Hussein, M., Elsebaie, E.H., Ali, S.H., Abdel Fattah, A.M., Hassan, M., Yassin, A., Mahfouz, N.A., Elsayed, R.M., et al. (2023). Neuropsychiatric post-acute sequelae of COVID-19: prevalence, severity, and impact of vaccination. Eur Arch Psychiatry Clin Neurosci 273, 1349–1358. 10.1007/s00406-023-01557-2.

7. Shariff, S., Uwishema, O., Mizero, J., Devi Thambi, V., Nazir, A., Mahmoud, A., Kaushik, I., Khayat, S., Yusif Maigoro, A., Awde, S., et al. (2023). Long-term cognitive dysfunction after the COVID-19 pandemic: a narrative review. Annals of Medicine & Surgery 85, 5504–5510. 10.1097/MS9.0000000000001265.

8. Ross, E.C., Olivera, G.C., and Barragan, A. (2022). Early passage of Toxoplasma gondii across the blood–brain barrier. Trends in Parasitology 38, 450–461. 10.1016/j.pt.2022.02.003.

9. Le Guennec, L., Coureuil, M., Nassif, X., and Bourdoulous, S. (2020). Strategies used by bacterial pathogens to cross the blood–brain barrier. Cellular Microbiology 22. 10.1111/cmi.13132.

10. Leda, A.R., Bertrand, L., Andras, I.E., El-Hage, N., Nair, M., and Toborek, M. (2019). Selective Disruption of the Blood–Brain Barrier by Zika Virus. Front. Microbiol. 10, 2158. 10.3389/fmicb.2019.02158.

11. Huang, X., Hussain, B., and Chang, J. (2021). Peripheral inflammation and blood–brain barrier disruption: effects and mechanisms. CNS Neurosci Ther 27, 36–47. 10.1111/cns.13569.

12. Patabendige, A., and Janigro, D. (2023). The role of the blood–brain barrier during neurological disease and infection. Biochemical Society Transactions 51, 613–626. 10.1042/BST20220830.

13. Lee, M.-H., Perl, D.P., Nair, G., Li, W., Maric, D., Murray, H., Dodd, S.J., Koretsky, A.P., Watts, J.A., Cheung, V., et al. (2021). Microvascular Injury in the Brains of Patients with Covid-19. N Engl J Med 384, 481–483. 10.1056/NEJMc2033369.

14. Vogrig, A., Gigli, G.L., Bnà, C., and Morassi, M. (2021). Stroke in patients with COVID-19: Clinical and neuroimaging characteristics. Neuroscience Letters 743, 135564. 10.1016/j.neulet.2020.135564.

15. Lo Re, V., Dutcher, S.K., Connolly, J.G., Perez-Vilar, S., Carbonari, D.M., DeFor, T.A., Djibo, D.A., Harrington, L.B., Hou, L., Hennessy, S., et al. (2022). Association of COVID-19 vs Influenza With Risk of Arterial and Venous Thrombotic Events Among Hospitalized Patients. JAMA 328, 637. 10.1001/jama.2022.13072.

16. Daly, S.R., Nguyen, A.V., Zhang, Y., Feng, D., and Huang, J.H. (2021). The relationship between COVID-19 infection and intracranial hemorrhage: A systematic review. Brain Hemorrhages 2, 141–150. 10.1016/j.hest.2021.11.003.

17. Islam, M.A., Cavestro, C., Alam, S.S., Kundu, S., Kamal, M.A., and Reza, F. (2022). Encephalitis in Patients with COVID-19: A Systematic Evidence-Based Analysis. Cells 11, 2575. 10.3390/cells11162575.

18. Thompson, D., Brissette, C.A., and Watt, J.A. (2022). The choroid plexus and its role in the pathogenesis of neurological infections. Fluids Barriers CNS 19, 75. 10.1186/s12987-022-00372-6.

19. Ayub, M., Jin, H.K., and Bae, J. (2021). The blood cerebrospinal fluid barrier orchestrates immunosurveillance, immunoprotection, and immunopathology in the central nervous system. BMB Rep. 54, 196–202. 10.5483/BMBRep.2021.54.4.205.

20. Zhang, L., Zhou, L., Bao, L., Liu, J., Zhu, H., Lv, Q., Liu, R., Chen, W., Tong, W., Wei, Q., et al. (2021). SARS-CoV-2 crosses the blood–brain barrier accompanied with basement membrane disruption without tight junctions alteration. Sig Transduct Target Ther 6, 337. 10.1038/s41392-021-00719-9.

21. Pellegrini, L., Albecka, A., Mallery, D.L., Kellner, M.J., Paul, D., Carter, A.P., James, L.C., and Lancaster, M.A. (2020). SARS-CoV-2 Infects the Brain Choroid Plexus and Disrupts the Blood-CSF Barrier in Human Brain Organoids. Cell Stem Cell 27, 951–961.e5. 10.1016/j.stem.2020.10.001.

22. Bao, L., Deng, W., Huang, B., Gao, H., Liu, J., Ren, L., Wei, Q., Yu, P., Xu, Y., Qi, F., et al. (2020). The pathogenicity of SARS-CoV-2 in hACE2 transgenic mice. Nature 583, 830–833. 10.1038/s41586-020-2312-y.

23. Lu, H., Liu, Z., Deng, X., Chen, S., Zhou, R., Zhao, R., Parandaman, R., Thind, A., Henley, J., Tian, L., et al. (2023). Potent NKT cell ligands overcome SARS-CoV-2 immune evasion to mitigate viral pathogenesis in mouse models. PLoS Pathog 19, e1011240. 10.1371/journal.ppat.1011240.

24. Wu, Y., Zeng, J., Pluimer, B., Dong, S., Xie, X., Guo, X., Liang, X., Feng, S., Wu, H., Yan, Y., et al. (2020). Microvascular Injury in Mild Traumatic Brain Injury Accelerates Alzheimer-like Pathogenesis in Mice (Neuroscience) 10.1101/2020.04.12.036392.

25. Xie, X., Ma, G., Li, X., Zhao, J., Zhao, Z., and Zeng, J. (2023). Activation of innate immune cGAS-STING pathway contributes to Alzheimer’s pathogenesis in 5×FAD mice. Nat Aging 3, 202–212. 10.1038/s43587-022-00337-2.

26. Sun, S.-H., Chen, Q., Gu, H.-J., Yang, G., Wang, Y.-X., Huang, X.-Y., Liu, S.-S., Zhang, N.-N., Li, X.-F., Xiong, R., et al. (2020). A Mouse Model of SARS-CoV-2 Infection and Pathogenesis. Cell Host & Microbe 28, 124–133.e4. 10.1016/j.chom.2020.05.020.

27. Tietz, S., and Engelhardt, B. (2015). Brain barriers: Crosstalk between complex tight junctions and adherens junctions. Journal of Cell Biology 209, 493–506. 10.1083/jcb.201412147.

28. Armulik, A. (2005). Endothelial/Pericyte Interactions. Circulation Research 97, 512–523. 10.1161/01.RES.0000182903.16652.d7.

29. Arturi, F., Melegari, G., Giansante, A., Giuliani, E., Bertellini, E., and Barbieri, A. (2023). COVID-19 Biomarkers for Critically Ill Patients: A Compendium for the Physician. Neurology International 15, 881–895. 10.3390/neurolint15030056.

30. Lai, Y.-J., Liu, S.-H., Manachevakul, S., Lee, T.-A., Kuo, C.-T., and Bello, D. (2023). Biomarkers in long COVID-19: A systematic review. Front. Med. 10, 1085988. 10.3389/fmed.2023.1085988.

31. Bellon, M., Schweblin, C., Lambeng, N., Cherpillod, P., Vazquez, J., Lalive, P.H., Schibler, M., and Deffert, C. (2021). Cerebrospinal Fluid Features in Severe Acute Respiratory Syndrome Coronavirus 2 (SARS-CoV-2) Reverse Transcription Polymerase Chain Reaction (RT-PCR) Positive Patients. Clinical Infectious Diseases 73, e3102–e3105. 10.1093/cid/ciaa1165.

32. Esmailian, M., Vakili, Z., Nasr-Esfahani, M., Heydari, F., and Masoumi, B. D-dimer Levels in Predicting Severity of Infection and Outcome in Patients with COVID-19.

33. Siddiqui, R., Mungroo, M.R., and Khan, N.A. (2021). SARS-CoV-2 invasion of the central nervous: a brief review. Hospital Practice 49, 157–163. 10.1080/21548331.2021.1887677.

34. Stefely, J.A., Christensen, B.B., Gogakos, T., Cone Sullivan J.K., Montgomery, G.G., Barranco, J.P., and Van Cott, E.M. (2020). Marked factor V activity elevation in severe COVID -19 is associated with venous thromboembolism. American J Hematol 95, 1522–1530. 10.1002/ajh.25979.

35. Greene, C., Connolly, R., Brennan, D., Laffan, A., O’Keeffe, E., Zaporojan, L., Connolly, E., Cheallaigh, C.N., Conlon, N., Doherty, C., et al. (2023). Blood-brain barrier disruption in Long COVID-associated cognitive impairment (In Review) 10.21203/rs.3.rs-2069710/v2.

36. Krasemann, S., Haferkamp, U., Pfefferle, S., Woo, M.S., Heinrich, F., Schweizer, M., Appelt- Menzel A., Cubukova, A., Barenberg, J., Leu, J., et al. (2022). The blood-brain barrier is dysregulated in COVID-19 and serves as a CNS entry route for SARS-CoV-2. Stem Cell Reports 17, 307–320. 10.1016/j.stemcr.2021.12.011.

37. Wenzel, J., Lampe, J., Müller-Fielitz, H., Schuster, R., Zille, M., Müller, K., Krohn, M., Körbelin, J., Zhang, L., Özorhan, Ü., et al. (2021). The SARS-CoV-2 main protease Mpro causes microvascular brain pathology by cleaving NEMO in brain endothelial cells. Nat Neurosci 24, 1522–1533. 10.1038/s41593-021-00926-1.

38. Deffner, F., Scharr, M., Klingenstein, S., Klingenstein, M., Milazzo, A., Scherer, S., Wagner, A., Hirt, B., Mack, A.F., and Neckel, P.H. (2020). Histological Evidence for the Enteric Nervous System and the Choroid Plexus as Alternative Routes of Neuroinvasion by SARS-CoV2. Front Neuroanat 14, 596439. 10.3389/fnana.2020.596439.

39. Yang, A.C., Kern, F., Losada, P.M., Agam, M.R., Maat, C.A., Schmartz, G.P., Fehlmann, T., Stein, J.A., Schaum, N., Lee, D.P., et al. (2021). Dysregulation of brain and choroid plexus cell types in severe COVID-19. Nature 595, 565–571. 10.1038/s41586-021-03710-0.

40. Lee, M.H., Perl, D.P., Steiner, J., Pasternack, N., Li, W., Maric, D., Safavi, F., Horkayne-Szakaly, I., Jones, R., Stram, M.N., et al. (2022). Neurovascular injury with complement activation and inflammation in COVID-19. Brain 145, 2555–2568. 10.1093/brain/awac151.

41. Ramlall, V., Thangaraj, P.M., Meydan, C., Foox, J., Butler, D., Kim, J., May, B., De Freitas, J.K., Glicksberg, B.S., Mason, C.E., et al. (2020). Immune complement and coagulation dysfunction in adverse outcomes of SARS-CoV-2 infection. Nat Med 26, 1609–1615. 10.1038/s41591-020-1021-2.

42. Suzzi, S., Tsitsou-Kampeli, A., and Schwartz, M. (2023). The type I interferon antiviral response in the choroid plexus and the cognitive risk in COVID-19. Nat Immunol 24, 220–224. 10.1038/s41590-022-01410-z.

43. Schwabenland, M., Salié, H., Tanevski, J., Killmer, S., Lago, M.S., Schlaak, A.E., Mayer, L., Matschke, J., Püschel, K., Fitzek, A., et al. (2021). Deep spatial profiling of human COVID-19 brains reveals neuroinflammation with distinct microanatomical microglia-T-cell interactions. Immunity 54, 1594–1610.e11. 10.1016/j.immuni.2021.06.002.

